# Region-specific alterations of perineuronal net expression in postmortem autism brain tissue

**DOI:** 10.1101/2021.12.31.474655

**Authors:** Cheryl Brandenburg, Gene J. Blatt

## Abstract

Genetic variance in ASD is often associated with mechanisms that broadly fall into the category of neuroplasticity. Parvalbumin positive neurons and their surrounding perineuronal nets (PNNs) are important factors in critical period plasticity and have both been implicated in ASD. PNNs are found in high density within output structures of the cerebellum and basal ganglia, two regions that are densely connected to many other brain areas and have the potential to participate in the diverse array of symptoms present in an ASD diagnosis. The dentate nucleus and globus pallidus were therefore assessed for differences in PNN expression in human postmortem ASD brain tissue. While Purkinje cell loss is a consistent neuropathological finding in ASD, in this cohort, the Purkinje cell targets within the dentate nucleus did not show differences in number of cells with or without a PNN. However, the density of parvalbumin positive neurons with a PNN were significantly reduced in the globus pallidus internus and externus of ASD cases, which was not dependent on seizure status. It is unclear whether these alterations manifest during development or are a consequence of activity-dependent mechanisms that lead to altered network dynamics later in life.

## Introduction

The most recent estimate of autism spectrum disorder (ASD) prevalence rose to 1 in 44 children (Walensky et al., 2021), an 18% increase from the previous rate of 1 in 54 announced by the Centers for Disease Control and Prevention (Redfield et al., 2020). With this steady increase in ASD prevalence each year, it is becoming increasingly important to identify the underlying neurodevelopmental mechanisms that contribute to ASD symptoms. Although the brain regions involved and the molecular underpinnings implicated in ASD are diverse, a diagnosis centers on social communication and sensorimotor challenges within the domain of restricted and repetitive behaviors (APA, 2013). As many complex behaviors, such as language, require the integration of information from visual, auditory, tactile and motor systems, it has been proposed that people with ASD struggle to unify multisensory information into a single percept (Baum et al., 2015; Beker et al., 2018; Chan et al., 2016; Robertson & Baron-Cohen, 2017; Stevenson et al., 2016; Zhou et al., 2018). As such, brain plasticity mechanisms are implicated in ASD and have driven research into vulnerable aspects of critical period plasticity (Hansel, 2018; Reh et al., 2020; S. S.-H. Wang et al., 2014; Yousif Ismail et al., 2016) as well as its relationship to altered network dynamics and excitatory/inhibitory imbalance (Blatt et al., 2001; Gabard-Durnam et al., 2019; Gogolla et al., 2009; J. Hussman, 2001; Levin et al., 2017; Rubenstein & Merzenich, 2003).

The disruptions in critical period plasticity may lead to altered network dynamics later in life and have been connected to the function of parvalbumin (PV) positive interneurons (Reh et al., 2020). As recently reviewed in Ruden et al., 2021, PV interneurons are particularly vulnerable to a diverse range of stressors and have been linked to ASD. While the exact timing may differ between brain regions and cell-types, the closure of the critical period of plasticity coincides with the formation of perineuronal nets (PNNs) that surround the soma and proximal dendrites of PV interneurons (J.-H. Cabungcal et al., 2013; Fawcett et al., 2019; Reichelt et al., 2019).

PNNs are a specialized and condensed form of extracellular matrix (ECM), which are consistently found to have genetic susceptibility in ASD from genome wide association studies (GWAS) (Anney et al., 2010; J. P. Hussman et al., 2011; K. Wang et al., 2009; Weiss & Arking, 2009). Among these genes, Reelin, a disintegrin and metalloproteinase with thrombospondin motifs (ADAMTS) and semaphorins are some of the strongest evidence linking ASD to disruption of the ECM (for review see: Pantazopoulos & Berretta, 2016; Sorg et al., 2016). Additionally, studies of PNNs have been prominent in animal models of fragile X syndrome, where reduced PNN expression has been demonstrated in Fmr1 knockout mouse auditory cortex and amygdala with relevance to altered fear-associated memory (Reinhard et al., 2019). In another study, pharmacological inhibition or genetic reduction of matrix component metalloproteinase-9 results in increased PNN production surrounding PV neurons, which normalizes auditory deficits in Fmr1 knockout mice (Pirbhoy et al., 2020; Wen et al., 2018). Xia et al., 2021 utilized a valproic acid (VPA) mouse model of ASD and found differences in intensities of PV and PNN subpopulations that possibly contribute to the progression of ASD.

In contrast to animal model studies, there is a lack of literature on PNN distribution in postmortem samples from patients with idiopathic autism. Thus, the current investigation centered around regions of the brain with high PNN density and relevance to ASD-related behaviors. Postmortem brain samples were taken from both the dentate nucleus (DN) of the cerebellum that has high PNN expression (Blosa et al., 2016; Carulli et al., 2020; Hirono et al., 2021) as well as the globus pallidus (GP) of the basal ganglia (Adams et al., 2001; Jan-Harry Cabungcal et al., 2019). The DN and GP are sources of outgoing projections to the thalamic, motor, premotor and sensory cortices that can affect the functionality of excitatory cortical neurons and play critical roles in many ASD-related cognitive, sensory and motor behaviors. Therefore, PNN expression in these two critical regions was quantified in ASD compared to neurotypical samples. While the DN did not show differences in neuronal number or PNN expression, the GP had significantly reduced PNN expression in ASD cases that was not dependent on seizure status. Future studies may aim to clarify the role PNN and PV neurons may play in plasticity and, ultimately, an ASD diagnosis.

## Methods

### Postmortem Tissue

Human postmortem brain tissue was obtained from the University of Maryland Brain and Tissue Bank, a brain and tissue repository of the NIH Neurobiobank. ASD cases were confirmed through Autism Diagnostic Interview-Revised (ADI-R) scores and/or received a clinical diagnosis of ASD from a licensed psychiatrist, case demographics are provided in Table 1. The University of Maryland Brain and Tissue Bank (NIH Neurobiobank) is overseen by Institutional Review Board protocol number HM-HP-00042077 and de-identifies all cases before distribution to researchers.

**Table 1.** Postmortem tissue case demographics.

| Cases | GP | DN | Diagnosis | Age | PMI | Gender | Ethnicity | Cause of Death |
| --- | --- | --- | --- | --- | --- | --- | --- | --- |
| 914 | x | x | Control | 20 | 18 | M | Caucasian | Vehicle accident, multiple injuries |
| 1158 | x | x | Control | 16 | 15 | M | Caucasian | Cardiomegaly |
| 4337 | x | x | Control | 8 | 16 | M | African American | Blunt force neck injury |
| 4599 | x |  | Control | 23 | 18 | M | African American | Cardiac arrhythmia/anomalous coronary artery |
| 4787 | x | x | Control | 12 | 15 | M | African American | Asthma |
| 5030 | x |  | Control | 24 | 14 | M | African American | Reactive airway disease |
| 5113 | x | x | Control | 36 | 20 | M | African American | Pulmonary embolism |
| 5334 | x | x | Control | 12 | 15 | M | Hispanic | Hanging/suicide |
| 5376 | x | x | Control | 13 | 19 | M | Caucasian | Hanging |
| 5387 | x | x | Control | 12 | 13 | M | Caucasian | Drowning |
| 5566 | x | x | Control | 15 | 23 | F | African American | Hypertrophic cardiomyopathy |
| 5646 | x | x | Control | 20 | 23 | F | Caucasian | Reactive airway disease |
| 5669 | x | x | Control | 24 | 29 | F | African American | Hypertensive cardiovascular disease |
| 5705 | x | x | Control | 31 | 26 | M | Caucasian | Cardiac arrhythmia |
| 5759 | x | x | Control | 34 | 28 | M | Caucasian | Atherosclerotic cardiovascular disease |
| 5813 | x | x | Control | 20 | 24 | M | African American | Atherosclerotic cardiovascular disease |
| 5889 | x | x | Control | 27 | 12 | M | Caucasian | Acute pneumonia complicated by sepsis |
| 5893 | x | x | Control | 19 | 11 | M | Caucasian | Dilated cardiomegaly |
| 5922 | x | x | Control | 46 | 10 | M | Caucasian | Atherosclerotic cardiovascular disease |
| 5926 | x | x | Control | 21 | 27 | M | African American | Cardiac arrhythmia with probable sickle cell disease |
| 5958 | x | x | Control | 22 | 24 | M | African American | Dilated cardiomegaly |
| 3916 | x | x | Autism | 32 | 22 | M | Caucasian | Congestive heart failure |
| 4334 | x | x | Autism | 11 | 27 | M | Hispanic | Acute hemorrhagic tracheobronchitis |
| 4899 | x |  | Autism | 14 | 9 | M | Caucasian | Drowning |
| 5027 | x | x | Autism | 37 | 26 | M | African American | Obstruction of bowel due to adhesion |
| 5115* | x | x | Autism | 46 | 29 | M | Caucasian | Complications of pseudomyxoma peritonei |
| 5144 | x | x | Autism | 7 | 3 | M | Caucasian | Cancer |
| 5176 | x | x | Autism | 22 | 18 | M | African American | Subdural hemorrhage |
| 5278* | x |  | Autism | 15 | 13 | F | Caucasian | Drowning associated with seizure disorder |
| 5403 | x | x | Autism | 16 | 35 | M | Caucasian | Cardiac arrhythmia |
| 5419* | x | x | Autism | 19 | 22 | F | Caucasian | Natural/epilepsy |
| 5565* |  | x | Autism | 12 | 22 | M | African American | Seizure disorder, complications |
| 5574 | x |  | Autism | 23 | 14 | M | African American | Pneumonia |
| 5631 | x |  | Autism | 18 | 96 | M | Caucasian | Acute hepatic failure |
| 5771 | x | x | Autism | 27 | 5 | F | Caucasian | Undetermined |
| 5841 | x | x | Autism | 12 | 15 | M | Caucasian | Hanging |
| 5864* | x | x | Autism | 20 | 42 | M | Caucasian | Seizure disorder |
| 5878 | x | x | Autism | 27 | 42 | M | Caucasian | Peritonitis |
| 5939* | x | x | Autism | 21 | 22 | M | Caucasian | Cardiovascular related |
| 5940* | x | x | Autism | 29 | 20 | M | Caucasian | Epilepsy complicated by drowning |
| 5945* | x | x | Autism | 20 | 24 | M | Caucasian | Chronic pulmonary aspergillosis |
| 5978 | x | x | Autism | 11 | 21 | M | Caucasian | Smoke inhalation |
| 6033* |  | x | Autism | 14 | 25 | F | Caucasian | Seizure disorder |

\* At least one documented seizure

Nineteen control and 18 ASD formalin fixed age-, gender- and PMI-matched human DN blocks were dissected in a consistent anatomical location across cases. Similarly, blocks of GP were dissected so that both the GPe and the GPi were contained within the same block for a total of 21 control and 20 ASD cases. Blocks of DN and GP were chosen from the same cases where possible (Table 1). Blocks were rinsed, cryoprotected, flash frozen and stored at −80 °C until they were cut at 40 µm, in series, onto glass slides with a cryostat (Leica CM1950) and again frozen at −80 °C.

### Immunohistochemistry (IHC)

IHC was performed similarly to Hoffman et al., 2016. Five frozen DN sections (every sixth interval) on slides from each case were thawed, dipped in KPBS and dried on a slide drying rack before antigen retrieval in tris buffer (pH 9.0) in a preheated scientific microwave (Ted Pella) at 35°C, 150 Watts for 10 minutes. Sections were then placed in 1% hydrogen peroxide in KPBS for 20 minutes at room temperature. Three washes in KPBS at 35°C and 150 Watts for one minute were performed (all subsequent wash steps are performed in this manner). Non-specific blocking for 30 minutes in 8% horse serum in KPBS was completed before incubation in primary antibody (anti-HPLN1 1:150, R&D Systems 2608-HP) for 48 hours at 4°C. Sections were washed and placed in biotinylated secondary antibody (anti-goat 1:700, Vector Laboratories BA-9500) for one hour then washed again. After incubation in avidin-biotin complex (A/B) (Vector Laboratories PK-6100) for one hour, sections were rinsed first in KPBS then in 0.175 M sodium acetate before a 20 minute exposure to nickel (Sigma SIG-N4882) 3 3’-diaminobenzidine tetrahydrochloride hydrate (DAB) (Sigma SIG-32750) in sodium acetate. Sections were washed in sodium acetate and then KPBS before being dipped in distilled water and incubated in 1% neutral red (Sigma) for 30 minutes and subsequently run through a series of alcohol dehydrations. Sections were placed in xylene (SIG-534056) for eight minutes then mounted with DPX (Sigma SIG-06522) and coverslipped.

In the GP, all steps were the same, except after the first nickel DAB reaction with HPLN1, slides were washed then the primary step was repeated for 48 hours with anti-PV (1:300, Sigma P3088) and processed in the same manner with biotinylated anti-mouse secondary (1:600, Vector Laboratories BA-2000), but nickel was not included in the second DAB reaction to produce a brown reaction product instead of black. Neutral red was omitted on these sections.

### Imaging and quantification

A Zeiss Microbrightfield Stereoinvestigator system was used to quantify neuronal densities within the manually drawn contours. In the DN, contours were drawn just inside the border where neuron density was highest and excluded regions without strong staining, which created a ribbon-like outline (Figure 1). The density of neurons surrounded by a PNN, neurons without a PNN and total neuron numbers were estimated with the optical fractionator method then divided by the total estimated area using the Cavalieri method. The GP was quantified in a similar manner except that one circular contour contained the entire region of interest and neurons were only counted if they had both a PNN and a PV stained soma.

**Figure 1.**
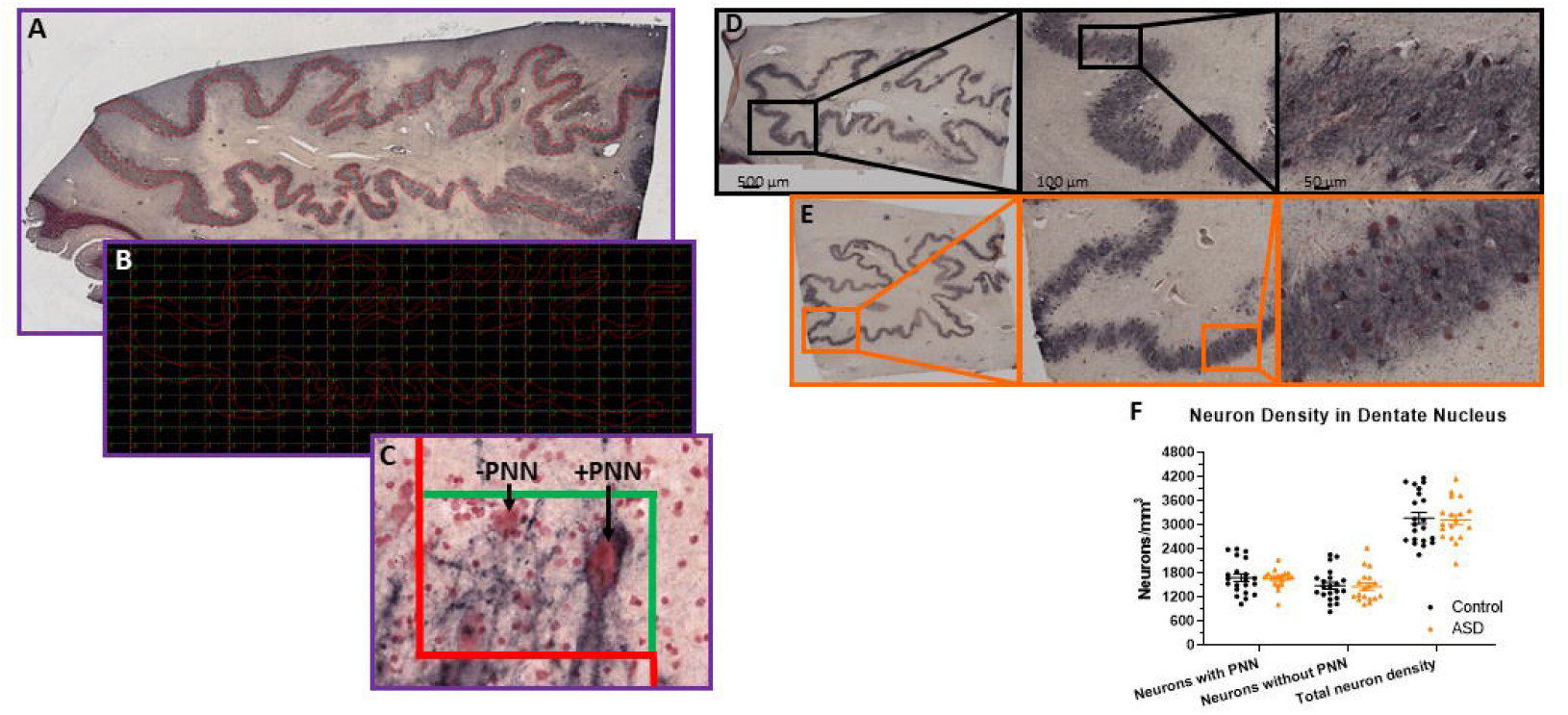
Stereological setup for quantifying perineuronal net density in human postmortem tissue. Contours were manually drawn over the region of interest, in this case (A), the dentate nucleus of the cerebellum. Contours were placed at a consistent distance from the edge of the dentate to mainly capture the high perineuronal net density locations in the center of the ribbon-like region. Therefore, the boxes actually counted within the grid placed over the contour in the software (B) would only be areas with a high density of neurons. Unstained regions are not represented. In this manner, unbiased counts can be achieved following established stereological methods, where neurons falling on the red line of the counting box (C) are not counted and any neurons falling within the box or on the green line are counted. Arrows point to examples of a dentate neuron surrounded by a perineuronal net and an example of a neuron stained with neutral red, but not surrounded by a perineuronal net. D) Immunohistochemistry of HPLN1 to visualize neurons surrounded by a perineuronal net while all neurons are stained with neutral red in a control (D) and ASD (E) case of the dentate nucleus within the cerebellum. F) The density of neurons surrounded by a perineuronal net (PNN), without a PNN and therefore the total density of all neurons within the dentate were not significantly different between control and ASD cases.

The Stereoinvestigator software has standardized programs to input counting grid sizes for the desired counting box coverage and automatically calculate neuron density following stereological principles (Gundersen, 1986; West et al., 1991). The counting grid size was adjusted until the number of boxes counted averaged 100-200 in the DN and 350-450 in each region of the GP of five 40 µm sections in each case series (every 6^th^ interval), resulting in a dissector volume of 0.00095 mm^3^.

## Results

Given the reported PC reductions in ASD (Bailey et al., 1998; M. Bauman & Kemper, 1985; M. L. Bauman et al., 1995; Skefos et al., 2014; Whitney et al., 2008; Hampson & Blatt, 2015; Schumann & Nordahl, 2011), we aimed to determine whether there are further deficits in neuronal numbers within the cerebellar circuitry, as DN neurons are likely to be impacted by PC deficits. PNNs around the DN neurons were also quantified as a readout for disrupted activity from PCs using hyaluronan and proteoglycan link protein 1 (HPLN1). HPLN1 is a critical component of PNN formation and has been shown to affect critical period plasticity after knockout (Carulli et al., 2010). Following stereological principles for estimating number within a volume, five sections for each case had a contour manually drawn around each region of interest (Figure 1A-C). The counting grid was adjusted to produce an average of 150 counting boxes per case and the total number of neurons with or without a PNN were totaled then divided by the number of counting sites times the dissector volume (0.00095 mm^3^).

The density of neurons (Figure 1D-F) with PNNs, based on HPLN1 expression, was not different (control mean= 1,682.27±417.51 and autism mean= 1,658.57±228.93 neurons/mm^3^, p=0.83). The density of neurons without a PNN also showed no differences (control mean= 1,487.74±386.44 and autism mean= 1,462.14±384.31 neurons/ mm^3^, p=0.84) and therefore total neuron numbers were similar (control mean= 3,170.01±632.64 and autism mean= 3,120.71±520.12 neurons/ mm^3^, p=0.79).

The shapes of the GPi and GPe (Figure 2A) allowed for circling the entire region of interest with one contour, with counting boxes filling the center. The same grid box size was utilized as for the DN, except the number of counting sites averaged 370 for the GPi and 442 for the GPe across the five sections in series of every 6^th^ interval. In pilot studies, there were not a significant portion of neurons that were either PV positive or PNN positive only, therefore, only neurons with a clear PNN and PV filled in the center were counted. PNN stain could be seen throughout the tissue as black “lines” following neuronal processes, but only somas with PV staining were counted as separate neurons.

**Figure 2.**
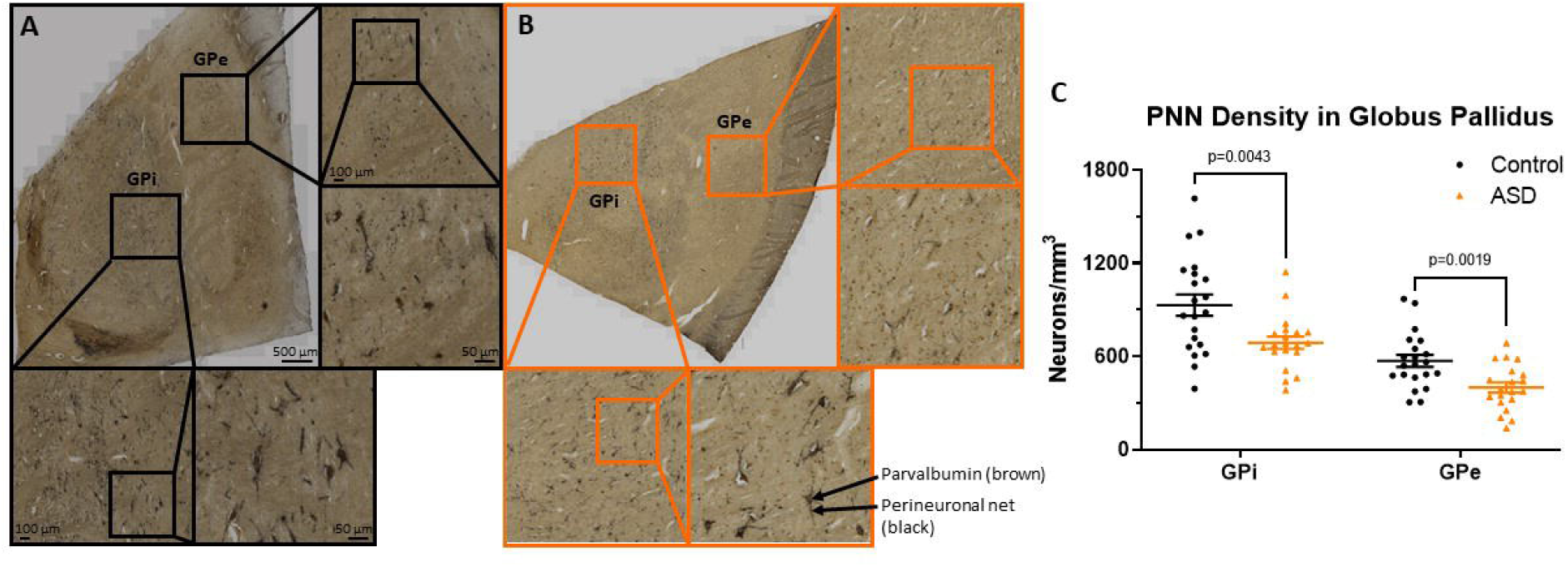
Immunohistochemistry of HPLN1 (black) and parvalbumin (brown) to visualize inhibitory projection neurons surrounded by a perineuronal in a control (A) and ASD (B) case of the globus pallidus (GP). The entire visible region of the GPi and GPe were each included in the contour manually drawn for inclusion within the counting grid. C) The density of parvalbumin positive neurons surrounded by a perineuronal net were significantly decreased in ASD cases compared to controls in both the GPi and GPe.

Neurons throughout the GPi and GPe that stained with both HPLN1 and PV were quantified and compared by diagnosis (Figure 2C), resulting in significant decreases in the ASD cases within the GPi (control mean 929.91±312.81 and autism mean 687.96±174.95, p=0.0043) and the GPe (control mean 570.46±178.85 and autism mean 399.54±146.89, p=0.0019).

In animal studies, degradation of PNNs occurs after seizures (Mcrae et al., 2012; Rankin-Gee et al., 2015; Yutsudo & Kitagawa, 2015) so the reduction of PNNs in the GP of ASD cases could be a result of seizure activity instead of a common feature across individuals. Within this cohort, 7/20 ASD cases also had a diagnosis of epilepsy, but comparing group means (Figure 3A) showed that the cases with seizures were not driving the decreases in lower PNNs for the ASD group as a whole. Instead, there was a trend toward higher PNN density in the seizure cases compared to the ASD only cases that did not reach significance (GPi p=0.24, GPe p=0.07).

**Figure 3.**
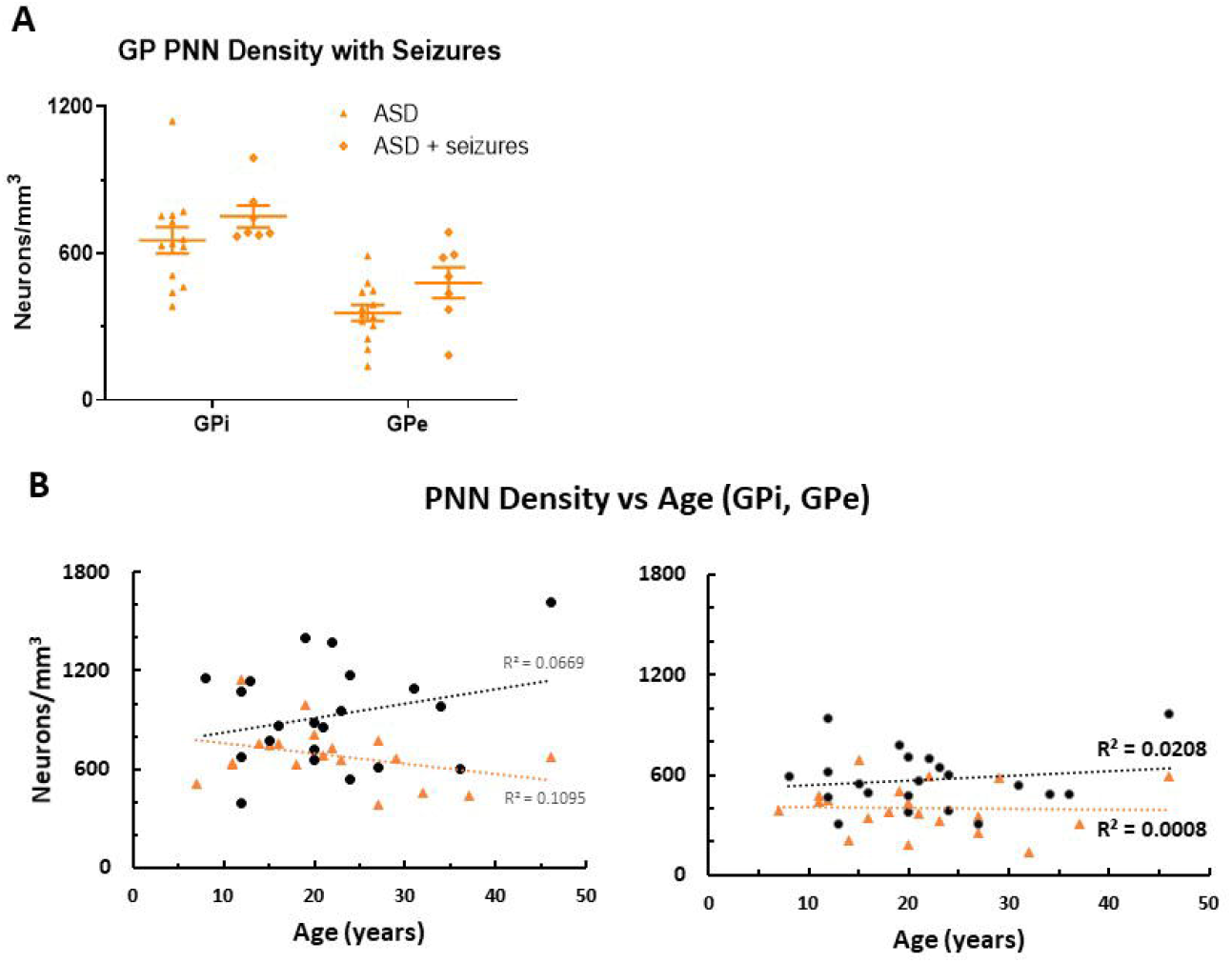
A) ASD cases were separated into groups based on medical reports of seizure occurrence. In the globus pallidus (GP), all seven seizure cases had a diagnosis of epilepsy. Although the group means were not statistically different, the seizure cases trended toward higher perineuronal net density than the ASD cases with no history of seizures. B) Perineuronal net density was plotted against age in both the GPi and GPe, which showed a positive correlation of density with age in the control group, but a negative correlation with age in the ASD group in the GPi, which was less pronounced in the GPe.

PNN density increases with age in human postmortem samples (Rogers et al., 2018). Therefore, PNN density was compared to age (Figure 3B), which showed a tendency to increase with age in controls, but was slightly decreased in ASD within the GPi. Cases had relatively the same density across age in the GPe.

Since the numbers of PNNs in ASD tissue were lower and the tissue often appeared to have a generally lighter stain (Figure 2), fresh frozen tissue from the opposite hemisphere from the same cases were assessed with western blotting in an attempt to quantify decreases in PNN component levels. However, these data were unreliable, as individual components of nets that should increase or decrease together, such as brevican and HPLN1, were highly variable even when measured from the same case on the same blot. Therefore, we did not include the results from these analyses, as discussed in the limitations section.

## Limitations

Although our number of cases included for comparisons is significantly higher than typically used in postmortem studies, the variability inherent to tissue processing and individual variation makes interpretation challenging. This is especially true for PNN analyses because, as other researchers have noted, variable fixation practices after brain collection, differences in collection strategies and the difficulties of extracting PNN components from surrounding tissue in a homogeneous manner all culminate in variance within the dataset (Rogers et al., 2018). In particular, even rigorous homogenization of tissue to release PNN components into solution proved difficult, as western blots were highly variable across cases and within a case for the different PNN component levels. As such, other strategies will need to be considered for quantifying amount of expression between cases. We quantified regions (DN, GPi and GPe) as a whole, so while we did not observe decreases in neuronal number within the DN, the results could be different if the area was subdivided into functional regions based on input from ASD affected areas in the cerebellar hemisphere. Since we did not have access to sections throughout the entire brain region of interest, true stereological assessments were not possible and will be useful for future studies when consistently fixed regions throughout an entire structure are available.

## Discussion

Despite reports of significant reductions in the number of PCs in the lateral hemisphere of the cerebellum, the targets of PC output within the DN appear to be sustained by the remaining PCs and inferior olive input. Individual neurons within the DN are each contacted by hundreds of PCs (Baumel et al., 2009; Chan-Palay, 1973) so it is reasonable that loss of a percentage of PCs would not impact numbers of DN neurons. The proportion of DN neurons surrounded by PNNs compared to those without appear to be unaltered as well. Future work may reveal differences in expression of PNN components that may be dependent on activity, but the variability inherent to current human postmortem methods did not allow for comparison of expression levels with IHC or western blots (see limitations section).

Within the GP, PNNs mainly surround PV positive projection neurons. This is in contrast to the DN neurons, in which PNNs mainly surround excitatory projection neurons that have input from PV positive PCs. Therefore, differences in ASD could be due to the different functional roles of these neurons subtypes. Both the GPi (with a higher percentage of PNNs) and the GPe had lower density counts in ASD tissue. It is typical for postmortem ASD studies to include seizure status as a covariate, as seizures can lead to many alterations and may be treated with medications that can affect the components being analyzed. One study measured PNN degradation and found that seizures lead to shifts in expression of PNN components, including HPLN1 (Mcrae et al., 2012), which may be due to release of matrix metalloproteinases following status epilepticus (Dubey et al., 2017; Dzwonek et al., 2004). However, a human postmortem study of PNN degradation after epilepsy did not show dysregulation of PNNs (Rogers et al., 2018), which may be dependent on the PNN markers examined and timing following seizures (reviewed in Chaunsali et al., 2021). Consequently, it is not clear whether PNNs degrade similarly in humans or whether the degradation is transient and potentially stabilizes over time.

Since the basal ganglia in general has been reported to have volume differences (for review: Subramanian et al., 2017) and receptor differences (Brandenburg et al., 2020) in ASD, the alterations in PNN density could be due to a general phenomenon of altered activity-dependent mechanisms. Emerging research implicates PV network dynamics in both critical period plasticity and adult learning, where experience related plasticity modulates learning and is dependent on a high or low PV network configuration (Donato et al., 2013; Hensch, 2005). Given that PNNs regulate input onto the soma of PV positive neurons, their digestion with chondroitinase ABC can alter this input and have effects on gamma oscillations, a key component of the role of PV neurons in network dynamics (Carceller et al., 2020). The development of complex skills, such as language and social communication, may be dependent on the function of PV neurons and PNNs in the temporal alignment of critical periods of plasticity across brain regions (Carulli & Verhaagen, 2021). Accumulating evidence implicates vulnerability of PV neurons in neuropsychiatric conditions (Ruden et al., 2021), making them intriguing targets for research and treatment strategies in ASD. The current study suggests alterations in PV neurons within the GP, which warrants further investigations into the function of these specialized neurons, as they are projection neurons that likely have different physiological functions compared to the typically studied PNN positive PV interneurons in cortical regions. As restricted and repetitive behaviors are core symptoms of an ASD diagnosis and the BG is a key regulator of repetitive behaviors, targeting these cell types could be valuable in mitigating undesirable symptoms.

## Acknowledgements

This work was supported by the Hussman Foundation.

The authors gratefully thank the families that donated brain tissue to the University of Maryland Brain and Tissue Bank (NIH NeuroBioBank) that make this and other research possible.

The authors are grateful for technical support from Charity Ensor, Abimbola Oladele and Brittany White.

